# The NuA4 acetyltransferase and non-canonical chromatin-associated MRN/ATM activity temporally define senescence-associated secretory programs

**DOI:** 10.1101/413732

**Authors:** Nicolas Malaquin, Audrey Carrier-Leclerc, Aurélie Martinez, Mireille Dessureault, Sabrina Ghadaouia, Jean-Philippe Coppé, Stéphanie Nadeau, Guillaume Cardin, Judith Campisi, Francis Rodier

## Abstract

**HIGHLIGHTS:**

- Tip60 complex cooperate with non-canonical DDR activity to trigger paracrine signals
- Low-level chromatin-associated ATM activity is required for the SASP
- The intensity and temporal dynamics of the SASP can be manipulated

**eTOC blurb:** A temporal ballet between DNA damage response ATM/MRN and NuA4/Tip60 complexes control senescence-associated paracrine signals that are important during aging and cancer therapy, suggesting that DNA damage responses, acetylation, and chromatin remodeling can be exploited to manipulate senescence and benefit health.

**SUMMARY:** Senescent cells display senescence-associated (SA) phenotypic programs such as proliferation arrest (SAPA) and secretory phenotype (SASP), which mediate their impact on tissue homeostasis. Curiously, senescence-inducing persistent DNA double-strand breaks (pDSBs) cause an immediate DNA damage response (DDR) and SAPA, but SASP requires days to develop, suggesting it requires additional molecular events. Here, we show that pDSBs provoke delayed recruitment of MRN/ATM and KAT5/TRRAP (NuA4/Tip60) complexes to global chromatin. This coincided with activating histone marks on up-regulated SASP genes, whereas depletion of these complexes compromised the SASP. Conversely, histone deacetylase inhibition triggered accelerated MRN/ATM/Tip60-dependent SASP without pDSBs, interlacing acetylation and non-canonical DNA damage-independent, low-level DDR activity in SASP maturation. DDR/acetylation regulation of SA programs is preserved in human cancer cells, suggesting novel targets for modifying treatment-induced SA phenotypes. We propose that delayed chromatin recruitment of acetyltransferases cooperates with non-canonical DDR signaling to ensure SASP activation only in the context of senescence, and not in response to a transient DNA damage-induced proliferation arrest.

## INTRODUCTION

Cellular senescence is a key tumor suppressor mechanism that relies on an irreversible senescence-associated (SA) proliferation arrest (SAPA) to limit the multiplication of cells at risk for neoplastic transformation (Lowe et al., 2004; Rodier and Campisi, 2011). Unlike apoptotic cells, which are rapidly eliminated (Christophorou et al., 2005; Christophorou et al., 2006; Roos and Kaina, 2013), senescent cells remain viable and have multifaceted biological functions, including roles in embryonic development (Munoz-Espin et al., 2013; Storer et al., 2013), viral infection (Appay et al., 2007), placental biology (Chuprin et al., 2013), wound healing (Demaria et al., 2014), and tissue remodeling that occurs during cancer treatment (Demaria et al., 2017; Gonzalez et al., 2015; Milanovic et al., 2018). Senescent cells are found with increased frequency in aging tissues and at sites of age-associated pathologies, including cancer, and the targeted elimination of senescent cells can restore naturally or prematurely aged tissue functions and extend lifespan in mice (Baker et al., 2016; Baker et al., 2011; Burd et al., 2013; Chang et al., 2016; Demaria et al., 2017; Dimri et al., 1995; Fuhrmann-Stroissnigg et al., 2017; Herbig et al., 2006; Jeon et al., 2017; van Deursen, 2014). While SAPA accounts for many tumor suppression functions of senescence, most of the detrimental, microenvironmental impact of senescent cells is mediated via their paracrine pro-inflammatory SA secretory phenotype (SASP) program (Alspach et al., 2014; Coppe et al., 2008; Krtolica et al., 2001; Laberge et al., 2015; Malaquin et al., 2016; Parrinello et al., 2005).

The establishment of SA phenotypes, in particular the SASP, requires a complex multiday genetic program that involves temporally interlaced molecular pathways (Baker and Sedivy, 2013; Ito et al., 2017; Rodier and Campisi, 2011). Senescence is almost always a consequence of genotoxic stresses (i.e., telomere shortening, oncogene-induced mitotic stress, irradiation) that cause DNA double-strand breaks (DSBs) and a DNA damage response (DDR) signaling cascade. In senescent cell nuclei, irreparable persistent DSBs (pDSBs) generate continuous DDR signaling from persistent nuclear structures termed ‘DNA segments with chromatin alterations reinforcing senescence’ (DNA-SCARS) (Fuhrmann-Stroissnigg et al., 2017; Fumagalli et al., 2012; Hewitt et al., 2012; Rodier et al., 2011). The DDR is necessary for both SAPA and SASP, but has important differences in temporal kinetics (Coppe et al., 2011; Kang et al., 2015; Malaquin et al., 2015; Pazolli et al., 2012; Rodier et al., 2009). For example, DSBs trigger immediate local recruitment of the MRE11-Rad50-NBS1 (MRN) complex on nearby chromatin, which promotes near-simultaneous ATM kinase recruitment/activation (characterized by the phosphorylation at serine 1981 (S1981-ATM)) that leads to DDR signaling and cell cycle checkpoints activation, including the CHK2 and p53/ p21^Cip1^ pathways controlling SAPA (Beausejour et al., 2003; Chen et al., 1995; d’Adda di Fagagna et al., 2003; Herbig et al., 2004; Rodier et al., 2009; Serrano et al., 1997). Alternatively, the DDR-dependent expression of many SASP factors is fully developed over a period of many days after the formation/presence? of senescence-inducing DNA lesions, long after the initial DDR signal has emerged from DSBs (Coppe et al., 2008; Coppe et al., 2011; Freund et al., 2011; Rodier et al., 2009; Rodier et al., 2011). Thus, despite its absolute requirement for initiation and maintenance of SASP, the canonical early activation of ATM and the DDR are not sufficient to trigger the SASP program, suggesting that non-canonical or partnered ATM activity could be involved (Rodier et al., 2009; Rodier et al., 2011).

DSBs are accompanied by a local reorganization of the chromatin important for the recruitment of the DNA repair machinery that accompanies emanating DDR signals. For example, the binding of the lysine acetyltransferase (KAT) complex Tip60 (enzyme KAT5) triggers local chromatin relaxation surrounding DSBs, contributing to ATM activation (Kaidi and Jackson, 2013; Squatrito et al., 2006; Sun et al., 2009). In fact, it is suggested that chromatin alterations themselves, even in absence of physical DNA lesion, can play a major role in the modulation of the DDR activity and perhaps in the establishment of SA phenotypes (Bakkenist and Kastan, 2003; Kaidi and Jackson, 2013; Soutoglou and Misteli, 2008). Similarly, earlier reports have shown that histone hyper-acetylation provoked by HDAC inhibitors (HDACi) triggers SAPA via the p53 and p16 pathways and is accompanied by SA-beta-galactosidase activity (SABGAL), another senescence hallmark (Munro et al., 2004; Ogryzko et al., 1996). HDACi-induced senescence was also shown to trigger the expression of IL-6 and IL-8 (Orjalo et al., 2009; Pazolli et al., 2012), but in this context, increased cytokine expression occurred in the absence of global γ-H2AX phosphorylation (Pazolli et al., 2012), suggesting that at least part of the DDR-associated SASP may occur in response to chromatin remodeling or “chromatin stress” rather than physical DNA breaks (Bakkenist and Kastan, 2003; Kaidi and Jackson, 2013; Pazolli et al., 2012).

Here, we compared SA secretory programs triggered by acetylation deregulation or direct DNA damage. We found that HDACi rapidly triggered an ATM-dependent SASP without DNA-SCARS, demonstrating that acetylation-associated chromatin stress can stimulate the DDR, independent of direct DNA damage. The SASP induced by HDACi treatment occurred rapidly within days compared to the SASP induced by direct DNA damage, which required up to a week to manifest. In both cases, global chromatin binding of chronic, low-level DDR-activated MRN/ATM and Tip60 complexes coincided with SASP development. Because HDACi are under investigation to treat multiple human cancers (West and Johnstone, 2014), this suggest that non-canonical DDR-mediated SASP induced by chromatin stress may contribute to the clinical outcomes of cancer therapy and may provide additional pharmaceutical targets to regulate SA phenotypes.

## RESULTS

### HDAC inhibition triggers a rapid SASP without classical DDR activation

To further investigate whether a canonical DDR is both essential and directly responsible for the SASP, we exposed normal human fibroblasts to the HDACi sodium butyrate (NaB), previously shown to trigger SAPA, SABGAL, and selected cytokine secretions without H2AX phosphorylation or obvious DNA damage (Ogryzko et al., 1996; Pazolli et al., 2012). Alternatively, we exposed cells to a single dose of 10 Gy X-Ray irradiation (XRA), which triggers a full senescence program associated with persistent DDR signaling and DNA-SCARS (Coppe et al., 2008; Rodier et al., 2009; Rodier et al., 2011). As expected, we immediately observed a stable proliferation arrest using either NaB or XRA (Fig 1a), and long-term treatment (9 days) generated SABGAL and IL-6 secretion as previously observed (Ogryzko et al., 1996; Pazolli et al., 2012) (Fig 1b, supplementary Fig S1a). We then probed whether sustained NaB treatment generated a SASP profile similar to that from typical DNA damage-induced senescence. The six most secreted proteins in replicatively senescent (REP) and irradiated (XRA) cells were also significantly increased by NaB (IL-6, GM-CSF, GRO, GRO-alpha, IL-8, and ICAM-1; *p* < 0.05), and overall, 17 proteins were differentially secreted at similar levels in both XRA and NaB-treated cells (*p* < 0.05) (Fig 1c). Importantly, factors that were secreted at lower levels after replicative senescence or XRA also decreased after NaB treatment, suggesting that HDACi does not trigger a global upregulation of secreted factors via non-specific chromatin deregulation but rather specifically activates SASP. Accordingly, the secretory profile (120 proteins) of NaB-treated cells correlated well with the profiles of cells that senesced by replicative exhaustion and irradiation, supporting common NaB-XRA SASP programs (supplementary Fig S1b). However, in contrast to irradiated cells where the SASP matures over a week or more (Coppe et al., 2008; Rodier et al., 2009), we noted that NaB-treated cells rapidly secreted high levels of IL-6 and IL-8. In fact, IL-6 and IL-8 secretion induced by NaB or by another HDACi (TrichostatinA (TSA)) was dose-dependent and detectable within 2 days after treatment (Fig 1d and supplementary Fig S1c-d), much earlier than secretion obtained following XRA, which reached similar levels after about 9 days (supplementary Fig S1a).

**Figure 1.**
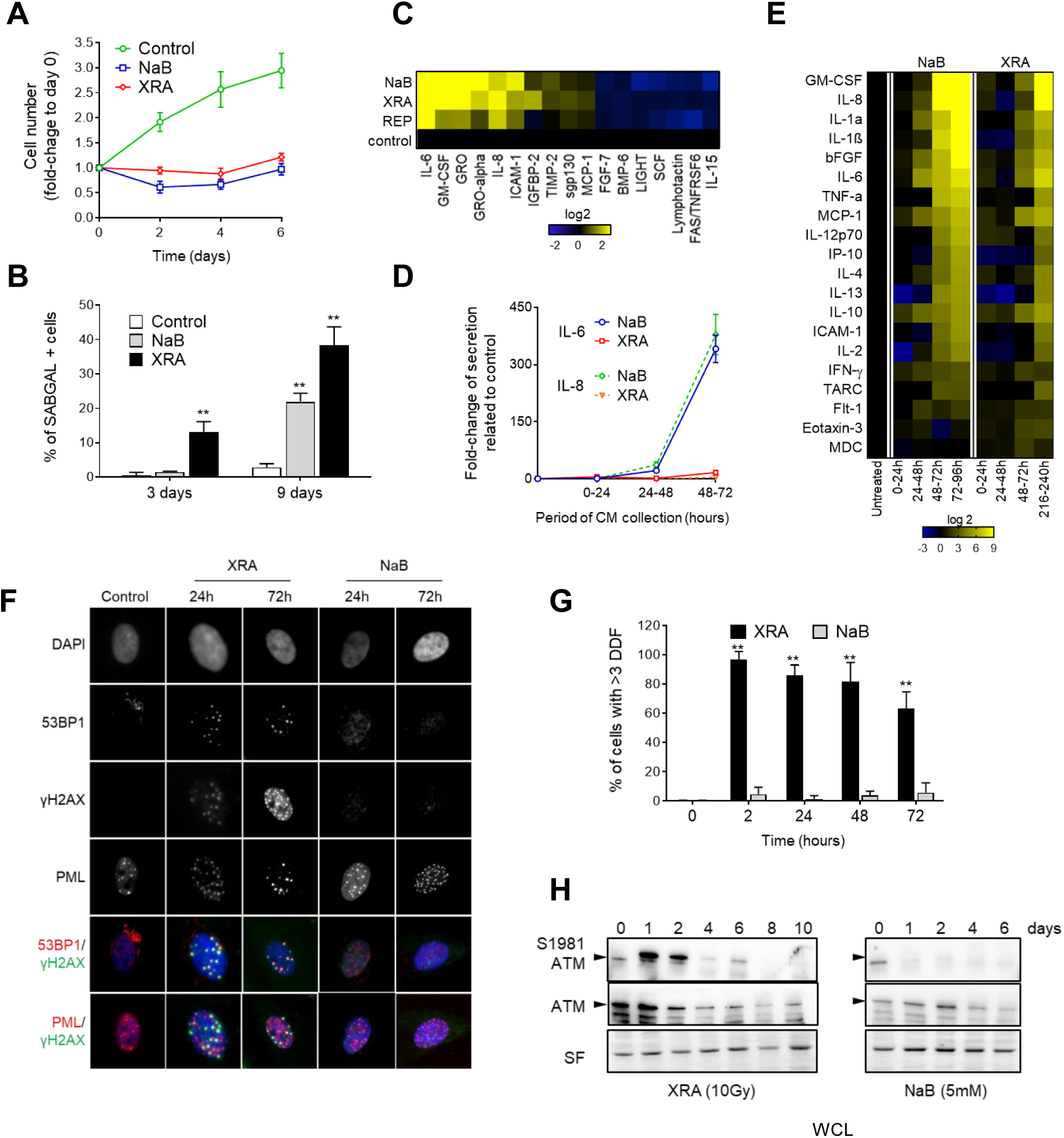
HDACi triggers a rapid DNA damage-independent SASP program. **(A**) Normal human fibroblasts (HCA2-hTert) were irradiated with 10 Gy X-rays (XRA) or treated with 5 mM sodium butyrate (NaB). Cells were fixed at 2, 4 and 6 days following treatment. Nuclear content was measured by fluorescent DNA dye (DRAQ5). For each condition, fluorescence intensity was reported on day 0. Data are mean ± S.D. and represent 3 independent experiments. **(B)** HCA2-hTert cells were untreated, exposed to 5 mM NaB or irradiated with 10 Gy. At indicated time, cells were fixed and assessed for the SA-ß-galactosidase assay. Data are mean ± S.D. and represent 3 independent experiments. **(C)** Soluble factors secreted by untreated or senescence-induced HCA2 cells [replicative senescence (REP); irradiation (10 days after 10 Gy of XRA)] or cells treated with NaB (2 mM for 10 days) were analyzed using antibody arrays. Only factors that were significantly modulated between control and treated (REP, XRA and NaB) at *p*<0.05 T-test are displayed. **(D)** HCA2-hTert cells were irradiated (10 Gy) or treated with NaB (5 mM). Serum-free conditioned media (SF-CM) were collected for the indicated periods, and IL-6 and IL-8 secretion was analyzed using ELISA. Data are mean ± S.D. and represent 3 independent experiments.**(E)** Soluble factors secreted by untreated (control), exposed to NaB (5 mM), or XRA (10 Gy) HCA2-hTert cells were analyzed for the indicated times by multiplex immunoassay (40-VPlex MSD). Data represent 2 independent experiments. For **(C)** and **(E)**, average secretion of control cells was a reference for baseline. Signals higher than baseline are yellow; signals below baseline are blue; unchanged fold variations are black (heat map key indicates log2-fold changes from the control). **(F)** Untreated, XRA and NaB-treated HCA2-hTert cells were fixed at indicated times and stained using immunofluorescence for 53BP1, γ-H2AX, PML nuclear bodies, and DAPI (nucleus counterstain). Top 4 rows display grayscale raw data for each staining channel. Bottom 2 rows display co-localization analysis with artificial color merges #1 and #2. Merge1 (top of the 2 rows?) displays DDF via colocalization (yellow) between 53BP1 (red) and γ-H2AX (green) with the nucleus in blue (DAPI). Merge2 (bottom of the 2 rows?) displays DNA-SCARS via colocalization (yellow) between PML nuclear bodies (purple) and γ-H2AX (green) with the nucleus in blue (DAPI). **(G)** Percentage of cells in (F) harboring 3 or more DDF (co-localization of γ-H2AX and 53BP1 in Merge1). Bars represent means ± S.D. from 3 independent measurements. T-test: ** *p*<0.01. **(H)** HCA2-hTert cells were irradiated with 10 Gy or treated with 5 mM NaB. Whole cell lysates prepared at indicated times were analyzed by western blotting. Data are representative of 3 independent experiments.

To further characterize the temporal kinetics of SASP factors induced by HDACi, we used sensitive multiplex immunoassays and compared the secretory profiles of 40 cytokines and chemokines from NaB-exposed cells (NaB-SASP) and irradiated cells (XRA-SASP). As expected, irradiation gradually increased secretion of pro-inflammatory factors including IL-6, IL-8, and GM-CSF, reaching SASP-levels in 10 days (Fig 1e, right panel). Alternatively, NaB triggered SASP-levels secretion or higher in only 3 days (Fig 1e, compare left and right panels). Levels of SASP factors that were elevated in the NaB-SASP at 24-48 h or 48-72 h were comparably lower or absent in corresponding XRA-SASP at all time points? (supplementary Fig 1e). Thus, HDACi strongly accelerate the expression of a mature SASP when compared to DNA damage.

Since SASP has been previously linked to persistent DNA lesions (Rodier et al., 2009; Rodier et al., 2011), we evaluated the presence of DSBs in the form of nuclear DNA damage foci (DDF) showing colocalization of 53BP1-γ-H2AX in NaB-and XRA-senescent cells. As expected, most XRA-treated cells rapidly (2 h) displayed 3 or more colocalized 53BP1-γ-H2AX DDF (Fig 1f) that transitioned to DNA-SCARS within 72 hours, as measured by gradual association of 53BP1-γ-H2AX DDF with promyelocytic leukemia (PML) nuclear bodies (Fig 1f) (Rodier et al., 2011). In contrast, NaB-treated cells did not display any increase in γ-H2AX foci, and the delayed formation of large 53BP1 foci that did not colocalize with γ-H2AX staining was not scored as DDF (Fig 1f-g). We then further probed DDR activation via the detection of ATM activating autophosphorylation at serine S1918 (Fig 1h), which was rapidly detected following exposure to irradiation (10 Gy), as expected for canonical DDR signaling, and then gradually decayed as cells repaired part of their DNA lesions and entered senescence (Fig 1h, left panel). However, ATM activation remained undetectable in NaB-treated cells even at several days after treatment initiation when cells have already developed a mature SASP profile, supporting the idea that DSBs are not generated and canonical DDR signaling is not activated in this context (Fig 1h, right panel). Likewise, the phosphorylation of the ATM target CHK2 (phospho T68-CHK2; (Ahn et al., 2000)) was rapidly observed after irradiation, but remained undetected following NaB exposure (supplementary Fig 1f). Taken together, these results suggest that the SASP does not always correlate with the formation of DSBs lesion or with the degree of typical (canonical) ATM/DDR activity detected, particularly in HDACi-induced senescent cells.

### ATM activity is required for HDACi-induced SASP

Because ATM-dependent DDR signaling is crucial for the production of oncogene-, XRA-, and REP-induced SASPs (Freund et al., 2011; Kang et al., 2015; Pazolli et al., 2012; Rodier et al., 2009; Rodier et al., 2011), we examined whether non-canonical (undetectable in whole cell lysate) ATM activity was nevertheless involved in NaB-induced SASP, despite the absence of DDF or DNA-SCARS. Using previously generated normal human fibroblasts that were infected with lentiviruses expressing short-hairpin RNA against green fluorescent protein (shGFP; control) or ATM (shATM) (Rodier et al., 2009), we observed that ATM depletion prevented IL-6 secretion triggered during NaB-induced senescence (Fig 2a). Similarly, NaB was not able to strongly enhance IL-6 secretion in primary A-T cells from patients carrying an inactivating mutation in the ATM gene, when compared to wild-type fibroblasts that usually show 100-fold increases in similar conditions (Fig 2b).

**Figure 2.**
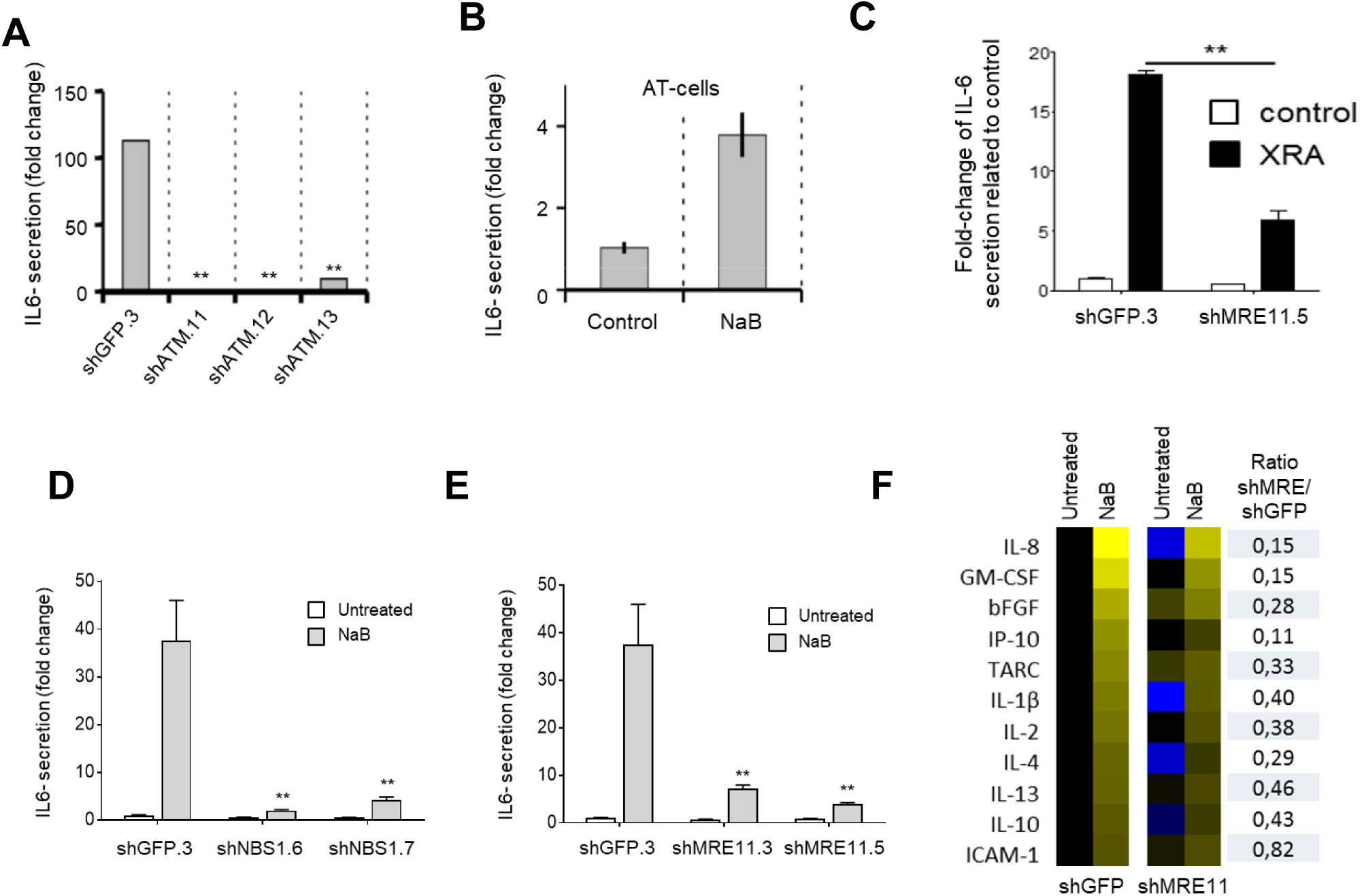
MRN and ATM trigger? the HDACi-induced SASP program. **(A)** HCA2 cells infected with lentiviruses expressing shGFP or shATM were allowed to recover for 7 days, and then treated for 2 days with 5 mM sodium butyrate (NaB). Conditioned media (CM) were collected over the next 24 h, and analyzed for IL-6 using ELISA. **(B)** A-T primary fibroblasts were either untreated or exposed to 2 mM NaB for 2 days. Serum free-CM (SF-CM) were collected over the next 24-hour interval for IL-6 ELISA. **(C-F)** BJ cells infected with lentiviruses expressing shGFP, shMRE11 or shNBS1 were allowed to recover for 7 days. **(C)** BJ-shGFP.3 and BJ-shMRE11.5 cells were untreated (control) or irradiated (XRA) with 10 Gy. After 9 days in culture, cells were put in serum-free medium for 24 hours. IL-6 secretion in SF-CM was analyzed by ELISA. Data are the means ± S.D. of triplicates and are representatives of 3 independent experiments. T-test: **p<0.01. **(D-E)** shRNA-expressing cells were treated for 2 days with 5 mM NaB. SF-CM were collected over the next 24 hours for IL-6 ELISA. Data are the mean ± S.D. of triplicates and represent at least 3 independent experiments. T-test: * *p*<0.05; ** *p*<0.01. **(F)** BJ-shGFP.3 and BJ-shMRE11.5 cells were either untreated or exposed to 5 mM NaB for 2 days, and CM were collected in serum-free conditions for the next 24 hours. Secreted soluble factors were evaluated using multiplex immunoassay (40-VPlex MSD). Average secretion of untreated BJ-shGFP.3 cells was used as baseline. Signals higher than baseline are yellow; signals below baseline are blue; unchanged fold variations are black (heat map key indicates log2-fold changes from control). For each soluble factor, the ratio of NaB-induced secretion between BJ-shMRE11 and BJ-shGFP was also calculated and displayed in the right column. Data represent 2 independent experiments.

To further evaluate the potential ATM activation process in the context of the HDACi-SASP, we depleted key components of the DNA damage sensor ATM-activating MRN complex, which is required for DNA damage-induced ATM activation and SASP (NBS1 (shNBS1) and MRE11 (shMRE11); supplementary Fig 2a) (Rodier et al., 2009; Shiloh and Ziv, 2013). In accordance with previous results for ATM and NBS1 (Rodier et al., 2009), irradiated MRE11-depleted cells showed decreased IL-6 secretion, confirming that the MRN complex is important for XRA-mediated SASP programs (Fig 2c). Surprisingly, the depletion of MRE11 or NBS1 prevented NaB-induced IL-6 secretion (Fig 2d and e), a finding validated using multiplex immunoassay profiling that revealed a systematic reduction in SASP factors following MRE11 depletion (Fig 2f). These results suggest that NaB-induced senescence occurs in the absence of DSBs DNA lesions, but nevertheless depends on the MRN/ATM DDR pathway, acting in a non-canonical way uncoupled from persistent DSBs-type DNA lesions. We thus confirmed that SASP-negative ATM-deficient cells could still form DDF after irradiation via ATR/DNAPK compensation mechanisms following the loss of ATM, as previously reported (supplementary Fig S2b) (Katyal et al., 2014). Overall, these results demonstrate that SASP correlates with low-level, hard to detect, non-canonical ATM activity rather than with DSBs DNA lesions themselves.

### HDACi triggers delayed chromatin recruitment of the ATM/MRN complex

NaB treatment does not provoke DSBs or canonical DDR activity (Fig 1f-h), but HDACi are known to non-specifically open chromatin structure via histone hyperacetylation (Jenuwein and Allis, 2001), suggesting a potential mechanism for SASP gene induction. However, the rapid chromatin hyperacetylation caused by HDACi, which occurs within a few hours of NaB exposure (Fig 3a), is not consistent with NaB-SASP program dynamics (Fig 1). More important, activating senescence via XRA leads to global histone hypoacetylation (Fig 3a), clearly suggesting that non-specific global histone hyperacetylation is not associated with SASP. Alternatively, following sustained NaB exposure, we observed the delayed formation of large 53BP1 foci that did not colocalize with γ-H2AX staining and correlated well with SASP kinetics (Fig 3b). This may have reflected higher level alterations of the chromatin structure reminiscent of 53BP1 bodies (Lukas et al., 2011). Since chromatin stress-mediated activation of the DDR was previously observed in the absence of DSBs (Kaidi and Jackson, 2013), we tested whether key SASP DDR mediators such as ATM and MRN could be detected directly on the chromatin rather than globally in the cell or within DDF. Using subcellular fractionation of NaB-and XRA-exposed cells that isolate and enrich for proteins from the soluble nucleus and chromatin compartment, we observed that low basal levels of soluble activated nuclear S1981-ATM gradually diminished in NaB-exposed cells, suggesting it may have relocalized (Fig 3c, left panel). Indeed, chromatin-associated fractions revealed that activated ATM was gradually recruited following NaB exposure, with kinetics corresponding to SASP maturation (Fig 3c, right panel). Likewise, the MRN complex component MRE11 was gradually recruited in chromatin fractions with similar kinetics, although it did not disappear from the nuclear soluble fractions (Fig 3c, right panel). In the case of XRA, we observed a strong increase of S1981-ATM in the nuclear soluble fraction early after exposure (4 h and 24 h, Fig 3d, left panel), confirming results obtained using whole cell lysate and consistent with high levels of XRA-induced DSBs at those moments (Fig 1h). However, S1981-ATM was strikingly undetectable in the global chromatin fraction. As cells repaired most DSBs (except DNA-SCARS (Rodier et al., 2011)), nuclear S1981-ATM gradually decreased over time and eventually returned to near basal levels 8-10 days post-XRA (Fig 3d, left panel). In contrast, S1981-ATM and MRE11 were slowly enriched in the global chromatin fraction, with kinetics similar to SASP maturation (Fig 3d, right panel), suggesting that chronic low-level recruitment of an activated MRN/ATM complex on the chromatin may temporally regulate the SASP program differently from high-level, nuclear and canonical DDR signaling observed immediately after DNA lesions.

**Figure 3.**
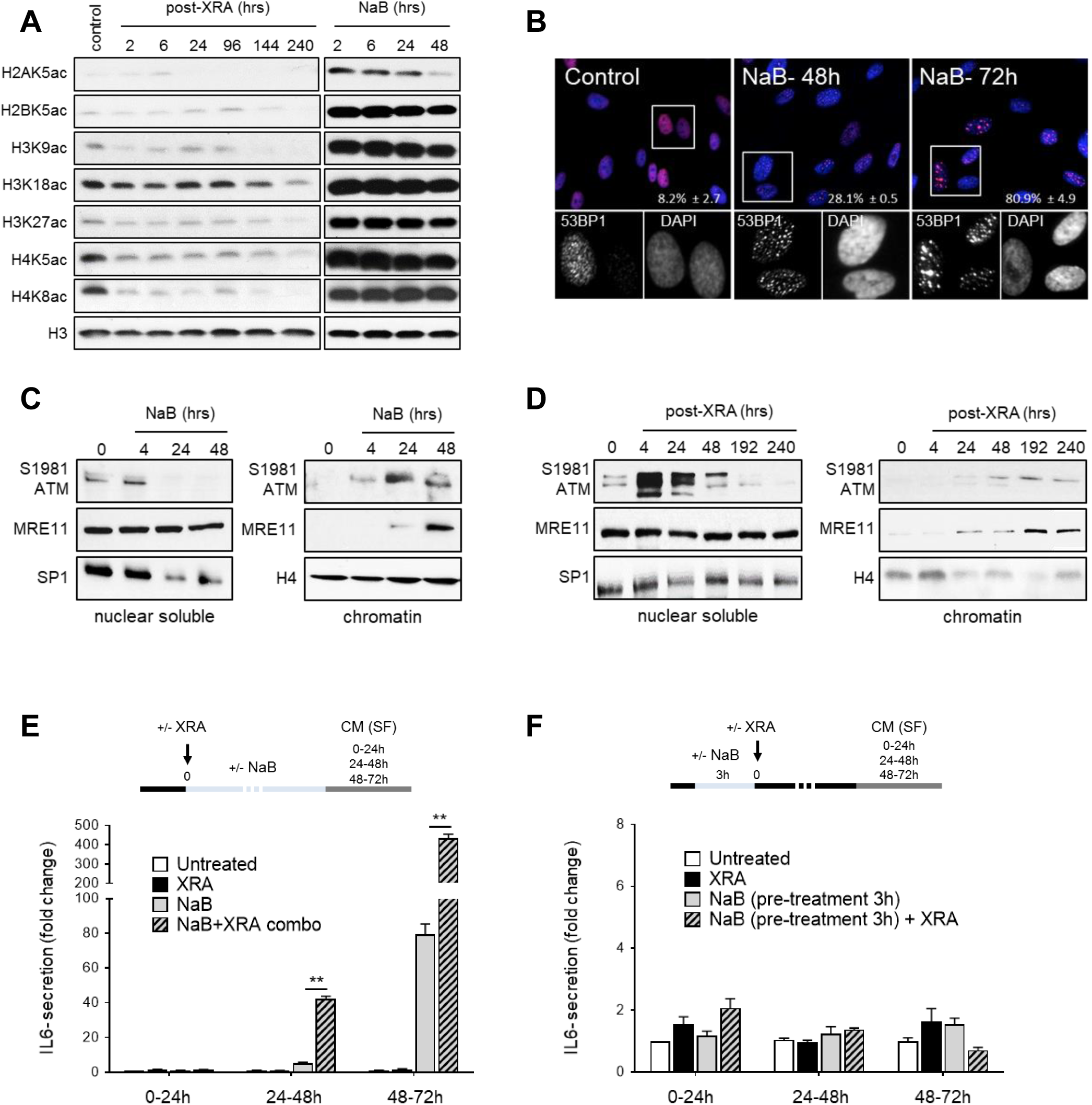
HDACi promote the binding of the MRE11/ATM complex to the altered chromatin. **(A)** HCA2-hTert cells were left untreated, irradiated with 10 Gy (XRA) or treated with 5 mM sodium butyrate (NaB). Whole cell lysates of indicated times were analyzed by western blot for histones or histone modifications. **(B)** HCA2-hTert cells were irradiated (10 Gy) or treated with 5 mM of NaB. At the indicated times, cells were fixed and stained for 53BP1 (red). Nucleus was stained with DAPI (blue). The percentage of cells in G that harbored 3 or more 53BP1 foci was calculated. Shown are the means ± S.D. from 3 independent measurements. **(C-D)** HCA2-hTert cells were **(C)** treated with 5 mM NaB or **(D)** irradiated with 10 Gy (time 0 = untreated controls). At the indicated times, cells were fractionated as described in the methods, and the nuclear soluble and chromatin fractions were used for western blot analysis of the indicated proteins. All results represent at least 3 independent experiments. **(E)** HCA2-hTert cells were left untreated, treated with 5 mM NaB alone, 10 Gy XRA alone or with a combination of both (NaB-XRA Combo). Serum-free conditioned media (SF-CM) containing 5 mM NaB were collected at indicated intervals for IL-6 analysis using ELISA. Data are means ± S.D. of triplicates and represent 5 independent experiments (\*\**p*<0.01). **(F)** HCA2-hTert cells were untreated or pre-treated with a pulse of NaB (5 mM) for 3 hours. NaB was removed and replaced by fresh medium. Cells were irradiated (or not) with 10 Gy XRA. SF-CM were collected at indicated intervals for IL-6 secretion. Data are reported as fold increase relative to untreated control cells. Data are means ± S.D. of triplicates and represent 3 independent experiments.

Because the HDACi-SASP programs depend on the slow chromatin accumulation of activated ATM/MRN, we reasoned that the availability levels of activated ATM could modulate this program. To test how HDACi-ATM signaling cooperatively activates the SASP, we simultaneously exposed normal fibroblasts to radiation (10 Gy) and NaB, and followed Il-6 secretion. While the first 24 hours post combo-treatment did not yield any significant increases in secretion, the cytokine was strongly detected after 24 hours when compared to either NaB or XRA alone (Fig 3e). Thus, DDR activity probably synergistically cooperates specifically with secondary HDACi molecular effects that take up to 24 hours to occur, excluding simple global histone hyperacetylation that occurs within a few hours. Furthermore, while a 2-hour NaB exposure was sufficient to trigger global histone hyperacetylation (Fig 3a), a similarly short pre-treatment with NaB (3 hours) just before XRA exposure was not sufficient to promote any synergistic increase in IL-6 secretion (Fig 3f).

### The Tip60 complex promotes NaB-and XRA-SASP

HDACi favor the activity of KAT complexes, resulting in increased histone and non-histone protein acetylation, chromatin relaxation, and promotion of gene expression (Davie, 2003; Stiehl et al., 2007). Because recent studies have associated the Tip60 complex to ATM activation during the initiation of the DDR induced by either DSBs or chromatin stress (Kaidi and Jackson, 2013; Sun et al., 2009), we hypothesized that Tip60 is a possible bridge between the XRA-and NaB-SASP programs. The depletion of Tip60 using two shRNA lentiviruses (shTip60) resulted in over 75% depletion compared to shGFP (Fig 4a) and had a negative impact on IL-6 secretion induced by NaB (Fig 4b), as well as most NaB-induced SASP factors including IL-8, MCP-1, and IL-1 (Fig 4c). Alternatively, the depletion of Tip60 (or MRN complex components) did not alter the SAPA induced by either NaB or XRA, demonstrating that different SA phenotypic programs can be functionally separated (supplementary Fig S3). To validate the involvement of the Tip60 complex, and not just the KAT5 protein, we depleted TRRAP (transactivation-transformation domain-associated protein), another component of the Tip60 complex important for chromatin binding (Robert et al., 2006). TRRAP depletion revealed a defect in IL-6 secretion induced by NaB when compared to control cells (Fig 4e), which extended to the secretion of many SASP factors (Fig 4f). Finally, the depletion of Tip60/TRRAP complex also negatively impacted IL-6 secretion induced by 10 Gy of XRA (Fig 4g), supporting a bidirectional cross-talk between DDR and KAT activities to regulate the SASP.

**Figure 4.**
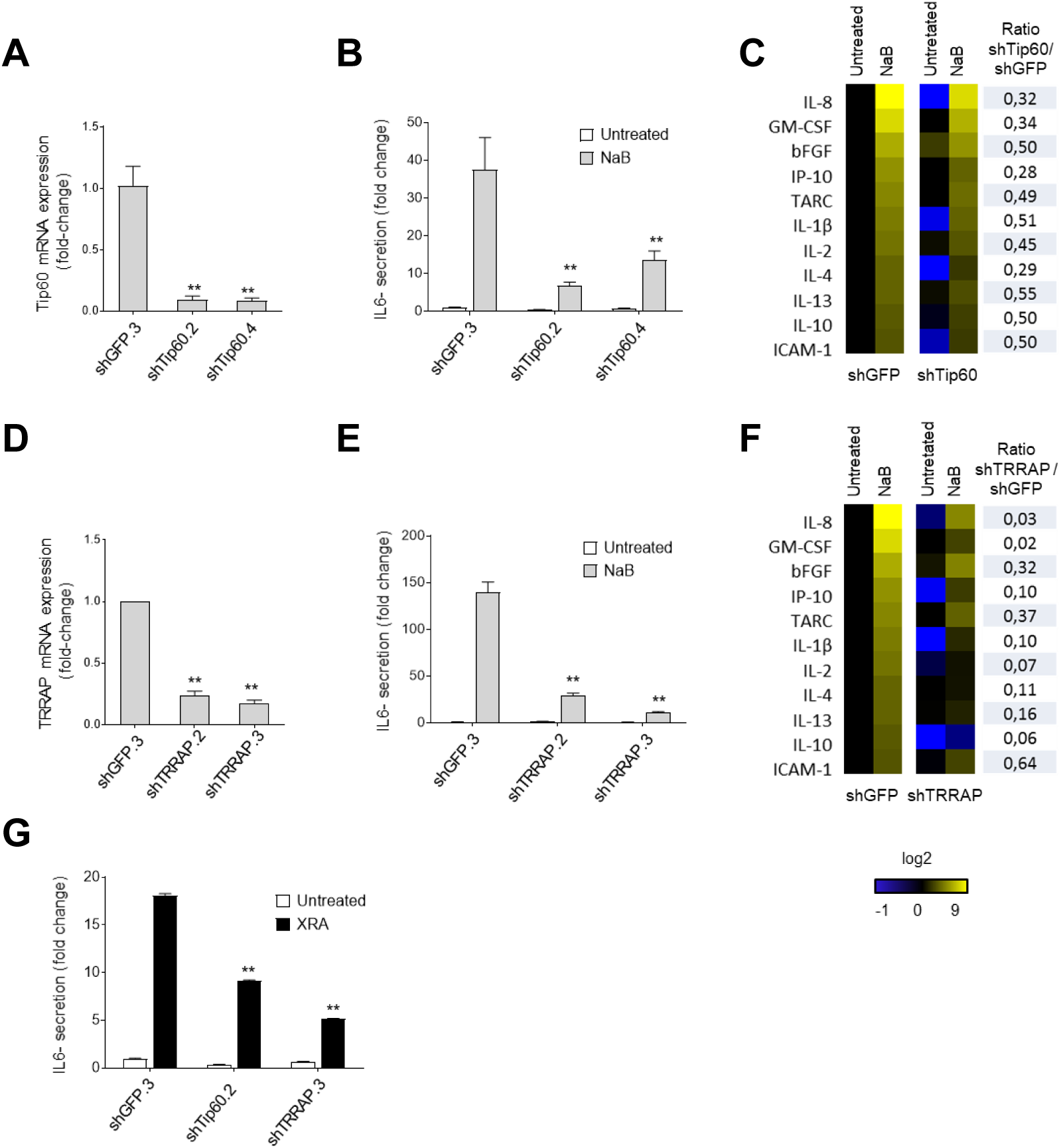
The Tip60/TRRAP acetyltransferase complex is essential for the HDACi-or XRA-induced SASPs. **(A)** BJ cells infected with lentiviruses expressing shGFP or shTip60 were allowed to recover for a minimum of 5 days. Expression of Tip60 mRNA in total RNA was analyzed by quantitative real time RT-qPCR (data normalized to 18S RNA). Bars represent the mean fold-change relative to shGFP cells in triplicate ± S.D. Data represent 2 independent experiments. **(B)** Cells in (A) were left untreated or treated for 2 days with 5 mM sodium butyrate (NaB). Serum-free conditioned media (SF-CM) were collected over the next 24 hours and analyzed for IL-6 using ELISA. **(C)** BJ-shGFP.3 and BJ-shTip60.2 cells were left untreated or exposed to 5 mM NaB for 2 days. SF-CM were collected for the next 24 hours. Secreted soluble factors were detected using multiplex immunoassay (40-Vplex MSD). **(D)** BJ cells infected with lentiviruses expressing shGFP or shTRRAP were allowed to recover for a minimum of 5 days. Expression of TRRAP mRNA in total RNA was analyzed and normalized as in (A). Bars represent the mean fold-change relative to shGFP cells in triplicate ± S.D. **(E)** Cells in (D) were left untreated or treated for 2 days with NaB (5 mM) and SF-CM were collected over the next 24 hours for IL-6 ELISA. **(F)** BJ-shGFP.3 and BJ-shTRRAP.3 cells were left untreated or exposed to 5 mM NaB for 2 days. SF-CM were collected for the next 24 hours. For (C) and (F), secreted soluble factors were detected using multiplex immunoassay (40-VPlex MSD). Average secretion of untreated BJ-shGFP.3 cells was used as baseline. Signals higher than baseline are yellow; signals below baseline are blue; unchanged fold variations are black (heat map key indicates log2-fold changes from control). For each soluble factor, the ratio of NaB-induced secretion between the BJ-shTip60 (or shTRRAP) and BJ-shGFP was calculated and displayed in the right column. Data represent 2 independent experiments. **(G)** BJ cells infected with shGFP.3, shTip60.2 and shTRRAP.3 were left untreated or irradiated with 10 Gy (XRA) and followed for 10 days. SF-CM were collected over the next 24 hours and analyzed for IL-6 ELISA. For ELISA analysis, data are reported as fold increase relative to untreated shGFP-cells. Data are means ± S.D. of triplicates and represent at least 3 independent experiments. T-Test: * *p*<0.05; ** *p*<0.01.

As endogenous Tip60 is notably difficult to detect, we tracked Tip60 movement within cell compartments in normal human fibroblasts using a lentivirus-delivered stably expressed HA tagged Tip60 (Tip60-HA) (Fig 5a), which did not impact basal or NaB-provoked levels of IL-6 secretion (Fig 5b). We then probed the kinetics of Tip60 movement in the soluble nucleus and chromatin compartments following NaB and XRA exposure. The recruitment of Tip60-HA on the chromatin was increased within 24 hours following NaB exposure (Fig 4c), while Tip60-HA recruitment was only observed beyond 96 hours after exposure to XRA (Fig 4d), in line with previously observed SASP kinetics under those conditions. The Tip60 complex can act as co-activator of NF-κB, the major transcription factor that regulates expression of SASP factors (Acosta et al., 2008; Kim et al., 2012; Salminen et al., 2012). Using subcellular fractionation, we observed that the NF-κB subunit p65 was increased in both the nuclear soluble and chromatin fractions of Tip60-HA cells treated with either NaB (48 h) or XRA (starting at 96 h) (Fig 5e), which was again consistent with SASP activation under those conditions. Similar increase of p65 level are also observed after NaB-treated or irradiated wild-type HCA2-hTert cells (supplementary Fig S4). To investigate more precisely if Tip60 was recruited directly on the promoters of SASP factors in relationship to NF-κB, we performed a Tip60 chromatin-immunoprecipitation (Ch-IP) to probe regions of NF-κB binding on IL-6 and IL-8 promoters. Tip60 Ch-IP revealed that NaB treatment (48 h) triggered an enrichment of binding on both IL-6 and IL-8 promoters when compared to an intronic region of RLP30 (negative control) (Fig 5f). This NaB-induced increase of Tip60 binding on Il-6 and IL-8 promoters was also accompanied by an elevation of histone H4 acetylation at lysine 8 (H4K8ac), a target of Tip60 (Squatrito et al., 2006), and of histone H3 acetylation at lysine 9 (H3K9ac), an epigenetic mark of an? active promoter (Fig 5f) (Guillemette et al., 2011). In response to DNA damage, Tip60 can directly interact with the MRN complex (Chailleux et al., 2010). Further probing the biochemical cross-talk between Tip60MRN complexes, we found that NaB treatment triggered co-immunoprecipitation of Tip60-HA with MRE11, suggesting that HDACi indeed promotes interactions between Tip60 and MRN complexes (Fig 5g).

**Figure 5.**
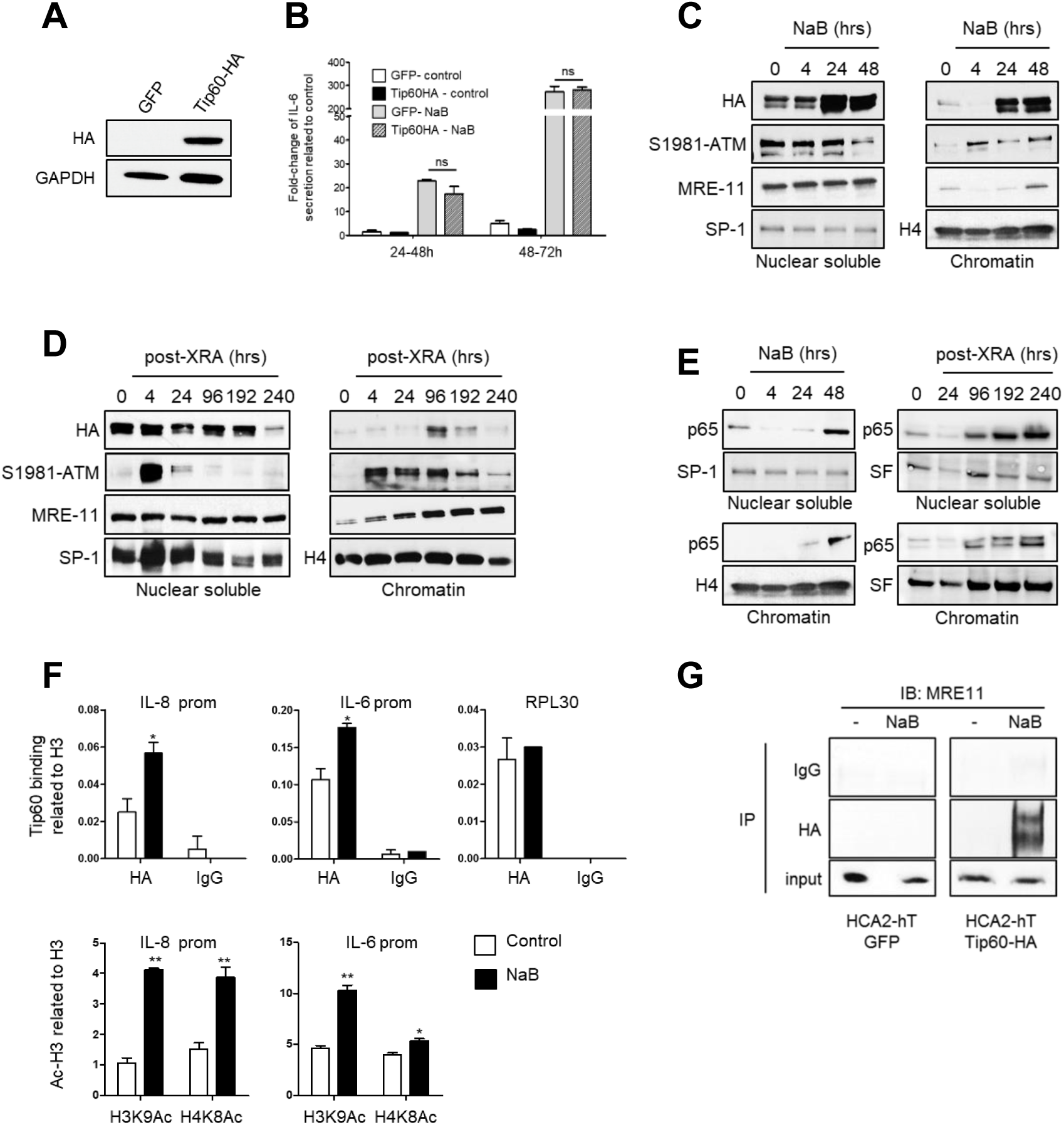
HDACi and XRA enhance the binding of Tip60 on SASP promoters. **(A)** HCA2-hTert cells were infected with lentiviruses expressing GFP or Tip60-HA and allowed to recover for 5 days. Whole cell extracts were collected and analyzed by western blotting for the indicated proteins. **(B)** Infected cells were untreated or treated with 5 mM of sodium butyrate (NaB) for 1 or 2 days, and the SF-CM were collected for the next 24 hours. IL-6 secretion was assessed using ELISA for the indicated 24-hour periods. The data are reported as fold increase relative to untreated control cells. Data are the means ± S.D. of triplicates and are representative of 3 independent experiments. T-test: *p<0.05; **p<0.01. **(C-E)** HCA2-hTert-Tip60-HA cells were treated with 5 mM NaB or irradiated with 10 Gy (XRA). At the indicated time, cells were fractionated as described in the methods and the nuclear soluble and chromatin fractions were used for western blot analysis of the indicated protein. All results represent at least 2 independent experiments. **(F)** HCA2-hTert-Tip60-HA cells were left untreated or treated with 5 mM NaB for 2 days. Chromatin was extracted and fragmented as described in methods. Chromatin immunoprecipitation (Ch-IP) was performed with indicated antibodies. Real-time qPCR was performed for the IL-6 and IL-8 promoters. Data, reported against histone? H3, are mean ± S.D. and represent 2 independent experiments. T-Test: * *p*<0.05; **
*p*<0.01. **(G)** HCA2-hTert cells infected with Tip60-HA or GFP were untreated or treated with 5 mM NaB for 2 days. Proteins were extracted for immunoprecipitation with HA or IgG antibodies. Precipitation of MRE11 was analyzed by western blot. Data represent 2 independent experiments.

### HDACi triggers senescence and SASP in prostate cancer cells

HDACi are emerging as a new class of chemotherapeutic agents for several types of cancers (West and Johnstone, 2014). They have been shown to induce tumor cell apoptosis, growth arrest, differentiation, and senescence, notably in prostate cancer (PCa) (West and Johnstone, 2014). To investigate whether HDACi-induced senescence and SASP are conserved in cancer cells, we exposed a metastatic PCa cell line (PC3) to XRA (control for senescence induction in PCa (Coppe et al., 2008)) or NaB, which both induced proliferation arrest and SABGAL in PC-3 cells (Fig 6a-b). Similarly to normal cells, NaB did not increased the number of 53BP1 DDF in PC-3 cells when compared to XRA, but generated 53BP1-body-like structures (Fig 6c). We observed the activation of a canonical DDR (phosphorylation of ATM and CHK2) after the XRA, which was virtually undetectable following NaB treatment (Fig 6d). Importantly, IL-6 secretion was increased 5 days after NaB or XRA exposure (Fig 6e). We then fully profiled PC-3 XRA-and NaB-SASPs and observed that most major SASP factors (IL-6, IL-8, IL-1, GM-CSF) identified in normal cells were also secreted (Fig 6g and Fig 1).

**Figure 6.**
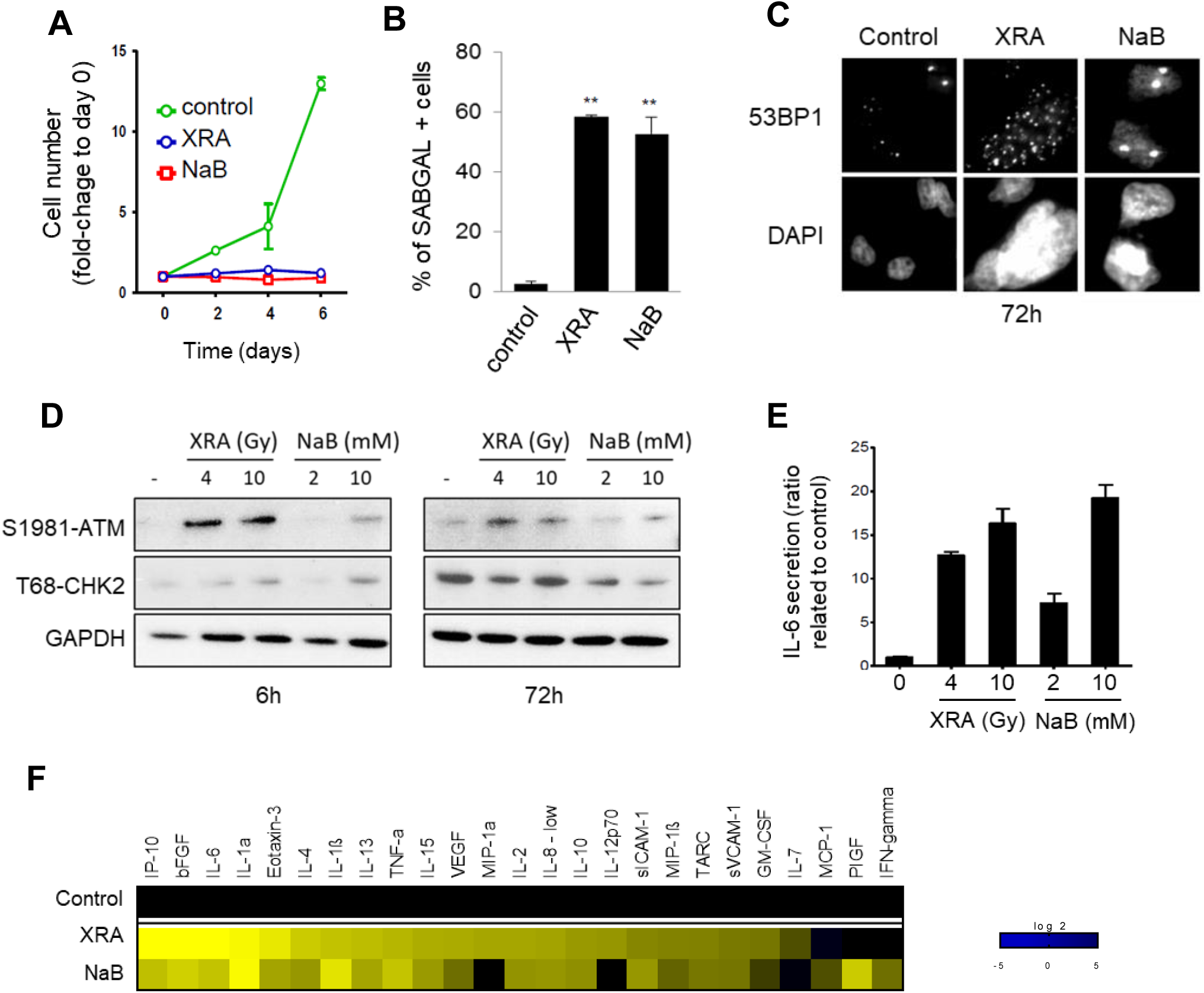
HDACi trigger senescence in prostate cancer cells. **(A)** Androgen-independent (PC-3) prostate cancer cell lines were untreated (control), exposed to 2 mM of sodium butyrate (NaB) or irradiated (XRA) with 4 Gy. Cells were fixed 2, 4 and 6 or 7 days after treatment and stained with the fluorescent DNA dye DRAQ5. Cell growth was evaluated using the fluorescence intensity. **(B)** Five days post-treatments of (A), SA-ß-Galactosidase assay were performed on PC-3. **(C)** PC-3 cells were untreated (control), exposed to 2 mM of NaB or irradiated with 10 Gy (XRA). At 72 hours later, the cells were fixed and processed for immunofluorescence detection of 53BP1 and nuclei were stained with DAPI. Representative images of 2 independent experiments are shown. **(D)** PC-3 cells were untreated (-) or treated with the indicated concentrations? of NaB or irradiated (XRA). At 6 and 72 hours later, whole cell extracts were collected and analyzed by western blotting for the indicated proteins. GAPDH was used as loading control. **(E)** PC-3 cells were treated for 5 days with NaB (2 and 10 mM) or irradiated with 4 and 10 Gy and cultured for 5 days. Serum-free conditioned media (SF-CM) were collected over the next 24 hours and analyzed for IL-6 by ELISA. The data are reported as fold increase relative to untreated control cells. **(F)** PC-3 were treated with NaB or irradiated as indicated. SF-CM were collected over the next 24 hours and analyzed for 40 soluble factors by multiplex immunoassay (40-VPlex MSD). Secretion of control cells was a reference for baseline. Signals higher than baseline are yellow; signals below baseline are blue; unchanged fold variations are black (heat map key indicates log2-fold changes from the control).

## DISCUSSION

SA phenotypic programs are essential for biological functions of senescent cells, and the underlying molecular mechanisms represent key targets for the manipulation of senescence in humans (Baker et al., 2016; Baker et al., 2011; Chang et al., 2016; Fuhrmann-Stroissnigg et al., 2017; Jeon et al., 2017; Zhu et al., 2017; Zhu et al., 2015). Among SA phenotypes, the microenvironment-remodeling SASP relies on a genetic program of interlaced intracellular and extracellular signaling pathways including DDR, p38MAPK, mTOR, the inflamasome, cytokines, and NOTCH (Acosta et al., 2013; Freund et al., 2011; Herranz et al., 2015; Hoare et al., 2016; Laberge et al., 2015; Malaquin et al., 2016; Rodier et al., 2009).

In addition to this multilayered molecular network, recent evidences confirm that SASP is a temporal multistep program involving distinct secretory profiles. In short, an early anti-inflammatory secretome regulated by NOTCH and rich in TGF-β is produced following the senescence stimuli and transiently counteracts the later establishment of a mature SASP containing more pro-inflammatory molecules such as IL-6 and IL-8, (Acosta et al., 2008; Hoare et al., 2016; Ito et al., 2017; Kang et al., 2015; Kuilman et al., 2008; Malaquin et al., 2015; Malaquin et al., 2016). We previously showed that the ATM-dependent DDR signalling is required for the establishment of a mature SASP as well as for the SAPA (d’Adda di Fagagna et al., 2003; Herbig et al., 2004; Rodier et al., 2009) and noted an unexplained disconnection between ATM activity levels and the SASP (Rodier et al., 2009). Indeed, the DDR is highly activated within minutes after inflicting DNA lesions, followed by proliferation arrest, whereas the DDR-dependent SASP takes days to develop. Here, we show a novel chronic, low-level, chromatin-associated, non-canonical ATM activity that is essential for the SASP program. This non-canonical ATM activity is apparently independent of physical DSBs DNA lesions or of the overall cellular levels of S1981 phosphorylated ATM. This is well illustrated in the SASP program induced by HDACi, which was ATM-dependent, yet associated specifically with the chromatin recruitment of ATM/MRN complexes and not with the presence of detectable DSBs. This suggests that for the SASP, association of an ATM complex with the chromatin is more important than the canonical DDR-associated, high-level ATM activation observed following the infliction of DNA lesions. We suggest that this is perhaps a mechanism to prevent SASP activation following simple DNA breaks that are easy to repair.

Our results show a key role for KAT5/Tip60 in the SASP program. Interestingly, we found a global decrease in KAT activities during senescence, as illustrated by a decrease of acetylated histone levels in senescent cells induced by XRA, consistent with previous studies reporting an alteration of histone acetylation patterns in aging tissues (Peleg et al., 2016). Alternatively, we found that rapid global chromatin hyper-acetylation caused by HDACi exposure was not sufficient in itself to trigger SASP, yet, HDACi still triggered a relatively rapid SASP program. This suggests that during the HDACi-SASP program, non-specific hyper-acetylation acts similarly to DNA damage, triggering a faster, but still multilayered non-canonical ATM/MRN activity at the chromatin that depends on the Tip60 complex. Despite the fact that global histone acetylation is decreased as cells enter senescence, we observed an enrichment of active epigenetic marks (i.e., H3K9ac) on SASP promoters, suggesting that specific KAT activities such as Tip60 are favored and essential to establish SA phenotypic programs. This supports the widespread idea that senescent cells reorganize the structure of their chromatin, and we now confirm that these cells present a specific pattern of acetylated regions in SASP factor genes (Narita et al., 2003; Shah et al., 2013). As acetylation is not restricted to histones, our data may also suggest that KATs involvement in SASP implicates the modulation of other non-histone proteins. KATs could control the expression of SASP factors by modulating activity of transcription factors like NF-κB, given that NF-κB acetylation is essential to regulate its diverse functions, including DNA binding activity, transcriptional activity or its ability to associate with IκBα (Chen et al., 2002; Huang et al., 2010). For example, Tip60 has been shown to modulate the transcriptional activity of RelA/p65 in NF-κB-dependent gene expression (Kim et al., 2012).

Importantly, the non-canonical DDR/KAT activities that we have identified to regulate SA phenotypic programs are conserved in cancer, at least for therapy-induced senescence in PCa, and suggests new pharmacological targets to manipulate senescence in the context of cancer therapy (Demaria et al., 2017). In the future, further understanding of the crosstalk between low-level, chromatin-associated, non-canonical DDR signaling and well-established SASP regulators including p38MAPK, mTOR, cytokine loops, and perhaps a greater acetyltransferase network, will help us decipher SA extracellular communication programs and may serve as a template to understand the regulation of other SA phenotypes like SAPA.

## ACKNOWLEDGEMENTS

We thank members of the Rodier laboratory as well as Christine Tam and Jacqueline Chung for valuable comments and discussions. We thank the microscopy & imaging platform of the Institut du cancer de Montréal. This work was supported by the Institut du Cancer de Montréal and by grants from the Canadian Institute for Health Research [MOP114962] and the Terry Fox Research Institute [1030] to F.R. F.R. is supported by a Fonds de Recherche Québec Santé junior I career award [22624]. N.M is supported by a Mitacs acceleration postdoctoral fellowship. A.CL., A.M., M.D., S.G., and S.N. are recipients of Institut du cancer de Montréal Canderel fellowships. S.N. is also an FRQS PhD fellowship recipient.

## AUTHORS CONTRIBUTIONS

N.M. and F.R. designed the study. N.M., A.CL., A.M., M.D., S.G., JP.C., S.N., and F.R. performed experiments and collected/analyzed data. G.C. provided technical assistance. N.M. and F.R. wrote the manuscript. F.R. provided financial support. J.C. provided technical support, expertise, conceptual advice, and revised the manuscript.

## MATERIAL AND METHODS

### Cell culture

HCA2 and BJ primary human foreskin fibroblasts were obtained from J. Smith (University of Texas, San Antonio) and cultured under ambient oxygen levels in Dulbecco’s modified Eagle’s media (DMEM) supplemented with 10% fetal bovine serum, 2.5 µg/ml fungizone and 100 U/ml penicillin/streptomycin. Unless specified, early passage fibroblasts were used and defined as having completed <35 cumulative population doublings (PD) and having a 24 h BrdU labeling index of >75%. PDs of primary cells were determined as follows: current PD = last PD + log2(cell number/cells seeded). Cell populations were considered replicatively senescent when cells achieved 24 h labeling indices of <5%. AT2SF primary A-T fibroblasts were obtained from the Coriell Institute and used at early passages (24 h BrdU labeling index of >75%). When indicated, the media was supplemented with sodium butyrate (Sigma) at 5 mM. The PC-3 cell lines were cultured in Roswell Park Memorial Institute medium (RPMI) supplemented with 10% fetal bovine serum, 2.5 µg/ml fungizone and 100 U/ml penicillin/streptomycin. The 293FT packaging cells (Invitrogen) were used to generate lentiviruses as previously described (Rodier 2011).

### DNA content-based cell counting

Cells were seeded in 96-well plates, treated as described, and fixed with formalin (Sigma) at indicated times. Relative cell quantity was determined by analyzing total DNA content using DRAQ5™ (Thermo Scientific) staining. Fluorescence intensity was measured using a Licor Odyssey scanner. Data is represented as a ratio of fluorescence intensity related to day 0.

### Live imaging-based cell proliferation assay

1500 cells/well were seeded in triplicate in 96-well plates, treated as described and incubated in the Incucyte Zoom System (Essen Bioscience). Cell number was imaged by phase contrast using the IncuCyte™ Live-Cell Imaging System (IncuCyte HD). Frames were captured at 2-hour intervals from two separate regions/well using a 10X objective. Proliferation growth curves were constructed using IncuCyte™ Zoom software, where plates were imaged and growth curves were built from the confluence mask.

### Viruses and infections

Viruses were produced as described previously (Campeau et al., 2009; Rodier et al., 2011) and titers adjusted when necessary to achieve ∼90% infectivity. Lentiviruses encoding shRNAs against GFP, ATM, MRE11, NBS1, TIP60, TRRAP, p300, CBP, GCN5 and PCAF were purchased from Open Biosystems. shRNA target sequences are provided in the supplemental material and methods. Tip60-HA and GFP were cloned into lentiviruses.

### Irradiation

Cells were X-irradiated with a total dose of 10 Gy at rates equal to or above 0.75 Gy/min using Gammacell® 3000 irradiator Elan.

### Senescence-associated β-galactosidase (SABGAL) detection

SAGAL assay was performed as previously described (Dimri et al., 1995). Briefly, cells were washed once with 1X PBS and fixed with 10% formalin for 5 min, then washed again with PBS and incubated at 37°C for 16 h in the staining solution containing at pH 6.0. The following day, cells were washed twice with PBS and pictures were taken.

### Immunofluorescence

Cells were fixed in Formalin for 10 min at room temperature and permeabilized in PBS-0.25% Triton for 10 min. Slides were blocked for 1 h in PBS containing 1% BSA and 4% normal donkey serum. Primary antibodies (listed in supplementary table 1) were diluted in blocking buffer and incubated with fixed cells overnight at 4° C. The cells were washed and incubated with secondary antibodies for 1 h at room temperature, then washed again and mounted with prolong + DAPI (Life technologies). Images were acquired on Zeiss fluorescence microscope with the axiovision software (Zeiss).

### Protein preparation and western blot analysis

Subcellular fractionation was performed using the subcellular protein fractionation kit (Thermo Scientific Pierce; # 78840) according to the manufacturer’s instructions. Whole cell lysates were prepared by scraping cells with mammalian protein extraction reagent (MPER, Pierce) containing protease and phosphatase inhibitors cocktail (Sigma). Protein concentration was measured using the bicinchoninic acid Protein Assay (BCA, Pierce). Proteins were separated on 7,5% or 4-15% gradient tris-glycine SDS-polyacrylamide gels (BioRad) and transferred onto PVDF membranes (Hybond-C Extra, Amersham). Primary antibodies used are listed in supplementary table 1. Primary antibodies were revealed with peroxidase-conjugated secondary antibodies (Santa cruz), followed by enhanced chemiluminescence (Pierce). Alternatively, secondary antibodies coupled with the indicated fluorescent dye (Licor) and fluorescence was detected using a Licor Odyssey infrared scanner. In this case, fluorescent bands were quantified using the software Image Studio™ Lite. Quantification from the GAPDH was used to normalize the results.

### SASP factors secretion measurement in conditioned medium

Conditioned media (CM) from control and senescent cells were prepared as previously described (Rodier, 2013). Briefly, CM was prepared by washing cells 3 times in serum-free medium, followed by incubation in serum-free DMEM containing antibiotics for 16 h. CM were collected and stored at −80°C until assayed. CM were assessed for ELISAs (Il-6 or IL-8) using kits and procedures from R&D (IL-6 #D06050, IL8 #DY208). The data was normalized to cell number and reported as fold change of secreted protein per cell per day in relation to the control. CM were also analyzed using antibody arrays (Chemicon, Human Arrays VI and VII, cat #AA1001CH-8) according to the manufacturer’s instructions with modifications as previously described(Coppe et al., 2008). Finally, CM were also analyzed using multispot electrochemiluminescence immunoassay system for 40 secreted factors using the V-Plex human kit from Meso Scale Discovery (MSD; #K15209D) following the manufacturer’s instructions.

### RT-qPCR

RNAs were isolated with the RNeasy kit (Quiagen) and stored at −80°C. Reverse transcription of 1 µg total RNAs were assessed using the Superscript III first-strand reverse transcriptase kit (Invitrogen; 200 units) according to the manufacturer instructions. Primers for PCR (see table S2) were designed with NCBI-Primer-BLAST software (http://blast.ncbi.nlm.nih.gov/) and qPCR protocols followed recommendations for the StepOnePlus™ real-time system (Applied Biosystem). PCR amplification products were revealed using KAPA SYBR®fast green fluorescence (ABl Prism™ kit, Kapa Biosystems). Data analysis was performed using the StepOne™ software (Applied biosystem). All target gene transcripts were normalized to GAPDH.

### Chromatin immunoprecipitation

Chromatin preparation, immunoprecipitation (ChIP), and analysis was performed as described by the manufacturer (SimpleChIP Enzymatic Chromatin IP Kit (magnetic beads) from Cell Signaling Technology). The chromatin was immunoprecipitated using normal immunoglobulin G (IgG; negative control; CST #2729), rabbit anti-histone H3 (CST, #4620), rabbit-anti HA (Y-11, Santa Cruz), rabbit anti-acetyl-Histone H3K9 (Lys9) (ab10812; AbCam) or rabbit anti-acetyl-Histone H4K8 (Lys8) (CST, #2594). Two percent of the supernatant fraction from the chromatin lacking primary antibody was saved as the “input sample.” The immunoprecipitated DNA was subjected to qPCR as describe above with primers amplifying promoter regions of the selected genes (see table S2). Values were quantified using the comparative CT method and normalized to H3.

## FIGURE LEGENDS-SUPPLEMENTARY

**Supplementary figure 1**

**(A)** HCA2-hTert cells were irradiated with 10 Gy (XRA) or exposed to 5 mM sodium butyrate (NaB). After 9 days in culture, cells were put in serum-free (SF) medium for 24 hours. IL-6 secretion in SF-conditioned media (SF-CM) was analyzed by ELISA. **(B)** Correlation between NaB and other senescence-induced secretory profiles analyzed in figure 1C. Correlations between NaB vs. XRA or NaB vs. REP were done using control as a baseline and are depicted as log2-fold changes. **(C)** HCA2 cells were untreated or treated 0, 1 or 2 days with the indicated doses of NaB or TSA and CM were collected the next 24 hours during which cells were maintained in the presence of the drug. IL-6 secretion was analyzed by ELISA. Data are reported as fold increase relative to untreated control cells. **(D)** Dose response of NaB on IL-6 (left panel) and IL-8 (right panel) secretion in SF-CM of HCA2-hTert cells 48 hours after the treatment. Data are the means ± S.D. of triplicates and are representative of 2 independent experiments. **(E)** Ratio of secretion of GM-CSF, IL-8, IL-1α, IL-1β, bFGF and IL-6 between NaB-treated and XRA cells at indicated time points after treatment (D and E). Data represent 2 independent experiments. **(F)** HCA2-hTert cells were irradiated with 10 Gy or treated with 5 mM NaB. Whole cell lysates prepared at indicated times were analyzed by western blotting.

**Supplementary figure 2**

**(A)** BJ cells infected with lentiviruses expressing shGFP, shMRE11 or shNBS1 were allowed to recover for 7 days. Whole cell lysates were collected and depletion of NBS1 and MRE11 proteins were validated by western-blot. (B) Primary A-T cells, or HCA2 cells infected with shGFP or shATM expressing lentiviruses were irradiated with 10 Gy. Cells were fixed at 2 hours or? 6 days later, and analyzed by immunofluorescence for 53BP1 (red). The nuclei were counterstained using DAPI (blue).

**Supplementary figure 3**

BJ cells were infected with lentiviruses expressing shGFP, shMRE11.5, shNBS1.6 or shTip60.2, and allowed to recover for a minimum of 5 days. Infected cells were (A) untreated, (B) treated with 5 mM of sodium butyrate (NaB) or (C) irradiated with 10 Gy X-rays (XRA). Cells were fixed at 2, 4 and 6 days following treatment, and the nuclear content was measured using a fluorescent DNA dye (DRAQ5). For each condition, the fluorescence intensity was reported on day 0. Data are mean ± S.D. and are representative of 3 independent experiments.

**Supplementary figure 4**

HCA2-hTert cells were treated with 5 mM NaB or irradiated with 10 Gy (XRA). At the indicated time, cells were fractionated as described in the methods and the nuclear soluble fractions were used for western blot analysis of the indicated protein. Results are representatives of 3 independent experiments.

